# A novel stress response pathway regulates rRNA biogenesis

**DOI:** 10.1101/2020.08.16.250183

**Authors:** Witold Szaflarski, Mateusz Sowiński, Marta Leśniczak, Sandeep Ojha, Anaïs Aulas, Dhwani Dave, Sulochan Malla, Paul Anderson, Pavel Ivanov, Shawn M. Lyons

## Abstract

Production of ribosomes is an energy-intensive process owing to the intricacy of these massive macromolecular machines. Each human ribosome contains 80 ribosomal proteins and four non-coding RNAs. Accurate assembly requires precise regulation of protein and RNA subunits. In response to stress, the integrated stress response (ISR) rapidly inhibits global translation. How rRNA is coordinately regulated with the rapid inhibition of ribosomal protein synthesis is not known. Here we show that stress specifically inhibits the first step of rRNA processing. Unprocessed rRNA is stored within the nucleolus, and, when stress resolves, it re-enters the ribosome biogenesis pathway. Retention of unprocessed rRNA within the nucleolus aids in the maintenance of this organelle. This response is independent of the ISR or inhibition of cellular translation but represents an independent stress-response pathway that we term Ribosome Biogenesis Stress Response (RiBiSR). Failure to coordinately regulate ribosomal protein translation and rRNA production results in nucleolar fragmentation. Our study unveils a novel stress response pathway that aims at conserving energy, preserving the nucleolus, and prevents further stress by regulation of rRNA processing.

## INTRODUCTION

Production of ribosomes is a major energetic and pro-growth task. Some estimates suggest that nearly 60% of the cell’s energetic costs are a result of ribosome production (Warner et al. 2001). The expenditure of such a large percentage of energetic reserves is a result of the engagement of all three nuclear RNA polymerases (Pol I: 18S, 28S, 5.8S rRNA, Pol II: ribosomal protein mRNA, Pol III: 5S rRNA), coordination of ribosomal protein mRNA translation, modification of rRNAs and assembly in the nucleolus. Compounding these energetic demands is the requirement for the accurate processing of the ∼13,000 nucleotide pre-rRNA (47S rRNA) into mature 18S, 5.8S, and 28S rRNAs. Again, estimates suggest that 60% of *de novo* RNA synthesis is rRNA. Maturation of the 47S rRNA requires multiple endonucleolytic and exonucleolytic processing events. The identity and processes that regulate some of these nucleases are yet to be identified. Further, these rRNAs are heavily chemically modified, making them second only to tRNAs in terms of the percentage of modified nucleotides. The efficient assembly of ribosomes requires precise regulation of rRNA transcription, processing, modification, and delivery of newly synthesized ribosomal proteins from the cytoplasm to the nucleolus.

Transcription and many of the steps in ribosome assembly takes place in a specialized membrane-less organelle known as the nucleolus. Maintenance of the nucleolar structure is driven, in part, by a liquid-liquid phase separation (LLPS) (Brangwynne et al. 2009). Two main factors responsible for this process are RNA and RNA-binding proteins that contain intrinsically disordered regions and low complexity sequences (IDR/LCS). Two major IDR-containing proteins in the nucleolus are fibrillarin (FBL) and nucleophosmin (NPM) (Feric et al. 2016; Mitrea et al. 2016; Yao et al. 2019). Additionally, electrostatic interactions target proteins containing Nucleolar Localization Signals (NoLS) to the nucleolus (Martin et al. 2015). The concentration of RNA processing and RNA modification enzymes in the nucleolus and other nuclear bodies is thought to increase the efficiency of processing and modification (Dundr and Misteli 2010). Increases in nucleolar size or changes in nucleolar morphology are linked to increased growth demands owing to the necessity for new ribosomes in driving protein synthesis (Montanaro et al. 2008).

The nucleolus has emerged as a significant stress-regulated organelle [Reviewed in (Boulon et al. 2010)]. In response to many stresses, such as heat shock (Liu et al. 1996), serum starvation (Chan et al. 1985), nucleotide deprivation (Grummt and Grummt 1976), and UV irradiation (Zatsepina et al. 1989), the nucleolar structure is fragmented and disrupted (Rubbi and Milner 2003). In most of these instances, nucleolar disruption is a result of the inhibition of rRNA transcription. Nucleolar structure can be disrupted when levels of newly synthesized RNA driving LLPS are lowered. In response to particular stresses (e.g., heat shock or acidosis), RNAs are transcribed by RNA polymerase II between rDNA genes in regions known as intergenic spacers (IGS), leading to the formation of reversible functional amyloids that may be cytoprotective (Audas et al. 2012; Audas et al. 2016; Lyons and Anderson 2016).

Adverse environmental conditions activate cellular stress response pathways. Diverse exogenous stresses, such as thermal stress, viral infection, oxidative stress or nutrient deprivation, elicit an equally diverse array of cellular responses. However, the underlying goal of each of these responses is to promote survival. A significant element of these pathways is to redirect energy from housekeeping and pro-growth activities towards cell survival strategies. Upregulation of a subset of genes (e.g., *ATF4*) promotes survival during stress. However, the expression of the vast majority of genes is downregulated, both transcriptionally and translationally.

Our previous work has shown that acute stress causes global translation inhibition following eIF2α phosphorylation, which results in the formation of stress granules (SGs), non-membranous liquid-liquid phase separations of untranslated mRNPs. The process by which this occurs is known as the “integrated stress response (ISR).” The translation of mRNAs encoding ribosomal proteins is preferentially inhibited and targeted to SGs upon eIF2α phosphorylation (Damgaard and Lykke-Andersen 2011). Additionally, the translation of ribosomal protein mRNAs is a primary target of the second major stress-response pathway that centers on the activity of mammalian target of rapamycin (mTOR) (Thoreen et al. 2012). Inactivation of mTOR preferentially inhibits the translation of ribosomal protein mRNAs. Additionally, the transcription of rRNA genes by RNA Pol I depends upon active mTOR (Mayer et al. 2004). Therefore, mTOR modulation directly regulates both the translation of ribosomal protein mRNAs and the transcription of rRNA. This coordinate regulation is important because cells must balance rRNA synthesis with *de novo* ribosome protein synthesis. Failure to do so results in nucleolar stress (Yang et al. 2018), which would only further compound the initial cellular insult. While the translation of ribosomal protein mRNAs is similarly affected by activation of the ISR, it is not known how this is coordinately regulated with rRNA biosynthesis. Further, the ISR is activated under acute stress conditions (typically <30 min) and rapidly inhibits translation. Thus, there is little leeway to regulate rRNA biosynthesis coordinately with the rapid shutoff of translation. We hypothesize that failure in such coordination results in a misallocation of energetic resources leading to further dysfunction.

Here, we show that stresses that induce eIF2α phosphorylation and SG formation also cause inhibition of rRNA synthesis. Rather than directly inhibiting rRNA transcription by targeting the RNA polymerase I or associated basal transcription factors, leading to the disruption of the nucleolus, as has been shown for mTOR-dependent stress responses, we show that the first step in 47S rRNA processing is inhibited. As the rate of rRNA transcription is intrinsically tied to the efficiency of rRNA processing (Schneider et al. 2007), failure to convert 47S rRNA into the next pre-RNA intermediate (45S rRNA) has the ultimate effect of repressing the rate of rDNA transcription. Therefore, rRNA production is “paused” rather than being inhibited. The unprocessed pre-rRNA is stored within the nucleolus until stress has resolved, at which point it can re-enter the ribosome biogenesis pathway.

Moreover, this mechanism allows for the maintenance of nucleolar structure during stress as RNA is a contributing factor in promoting LLPS in RNA granules (such as SGs or nucleolus). Retention of unprocessed rRNA within the nucleolus during stress aids in the maintenance of nucleolar structure, such that when stress has passed, ribosome biogenesis machinery remains localized to the nucleolus. In contrast, direct inhibition of Pol I transcription compromises nucleolar integrity, thereby necessitating reassembly of the nucleolus after stress has passed. Further, we show that despite being coordinated with eIF2α phosphorylation, stress-responsive modulation of rRNA processing is regulated by an independent yet parallel signaling pathway. Finally, we show that failure to regulate rRNA production coordinately with translation results in nucleolar dysmorphology. Our data unveil a novel mechanism by which the first processing event in rRNA processing is regulated in a stress-dependent manner to conserve energetic reserves and maintains nucleolar structure during stress.

## Results

We began our investigation of the connection between rRNA biosynthesis and stress response by analyzing the effect of stress on rRNA transcription using the 5-ethenyl uridine (5-EU) CLICK-IT assay. U2OS osteosarcoma cells were stressed for 90 minutes with NaAsO_2_, the most widely used inducer of ISR, then 5-EU was added for an additional 30 minutes along with NaAsO_2_ (**Figure 1A**). Alternatively, cells were left untreated or were treated with Actinomycin D (ActD), a potent inhibitor of transcription. Cells were fixed, permeabilized, and Alexa488 fluorophore was conjugated to the incorporated 5-EU to visualize nascent RNA. To mark the nucleolus, we used antibodies against NPM, a resident of the granular component (GC) subcomponent of the nucleolus. In control conditions, a high concentration of nascent RNA is present in the nucleolus consistent with rRNA accounting for the vast majority of transcription (**Figure 1Ba**). We did not detect active transcription in the nucleolus after 90 minutes of ActD or NaAsO_2_ treatment by this assay (**Figure 1Bb – c**). However, this presented a conundrum since nucleolar integrity was disrupted by ActD but not NaAsO_2_. Therefore, we more directly interrogated rRNA by northern blotting. In humans, the primary ribosomal RNA transcript, termed the 47S rRNA, is polycistronic, containing the 18S, 28S and 5.8S rRNA. The mature rRNAs are flanked by 5’ and 3’ external transcribed spacers (5’ and 3’ETS) and separated by two internal transcribed spacers (ITS1 and ITS2). 47S rRNA is transcribed by RNA polymerase I and must be reiteratively processed by multiple enzymes to release the mature rRNAs (**Figure 1C**). The 5S rRNA is transcribed from other loci by RNA polymerase III. Using probes to different parts of the 47S transcript, we can analyze different rRNA processing intermediates. We began by analyzing the full 47S rRNA with a probe in the 5’ETS at the extreme 5’ end of the rRNA. Our previous work, and those of others, has shown that certain chemotherapeutic drugs are potent inducers of the ISR and a prevalent hypothesis has been that disrupting ribosome biogenesis would be a potent strategy in combating tumor progression (Szaflarski et al. 2016; Catez et al. 2019). Therefore, we expanded our analysis to include the chemotherapeutic drug lomustine. Similar to NaAsO_2_, lomustine induces phosphorylation of eIF2α, the hallmark of ISR induction, while ActD does not (**Figure S1A)**. However, both NaAsO_2_ and Lomustine treatments resulted in a striking increase in 47S precursor rRNA and the formation of a faster migrating species (**Figure 1D**, ln 3, 4). The changes in 47S expression were independently confirmed by qRT-PCR (**Figure 1E**). The faster migrating species that has alternatively been termed the 30S_+01_ fragment or the 34S fragment is often seen when small subunit (SSU) processome is inhibited, e.g., under RNAi mediated knockdown of certain RNA processing factors such as fibrillarin (Tafforeau et al. 2013). This fragment results from a failure to process the first cleavage site, the A’/01 site (**Figure 1C**), and spurious processing at the “site 2” site. Further northern blotting for ITS1 and ITS2 revealed a decrease in downstream processing intermediates (e.g. 41S, 26S, 21S, 18S-E and 12S) (**Figure 1D** and **S1B – E**). It is worth noting that northern blotting to ITS1 does not distinguish between the aberrant 34S product and canonical 30S precursors. Regardless, these data suggest that pre-rRNA processing is inhibited by preventing A’/01 processing, leading to an increase in 47S pre-rRNA and a decrease in downstream precursors.

**Figure 1.**
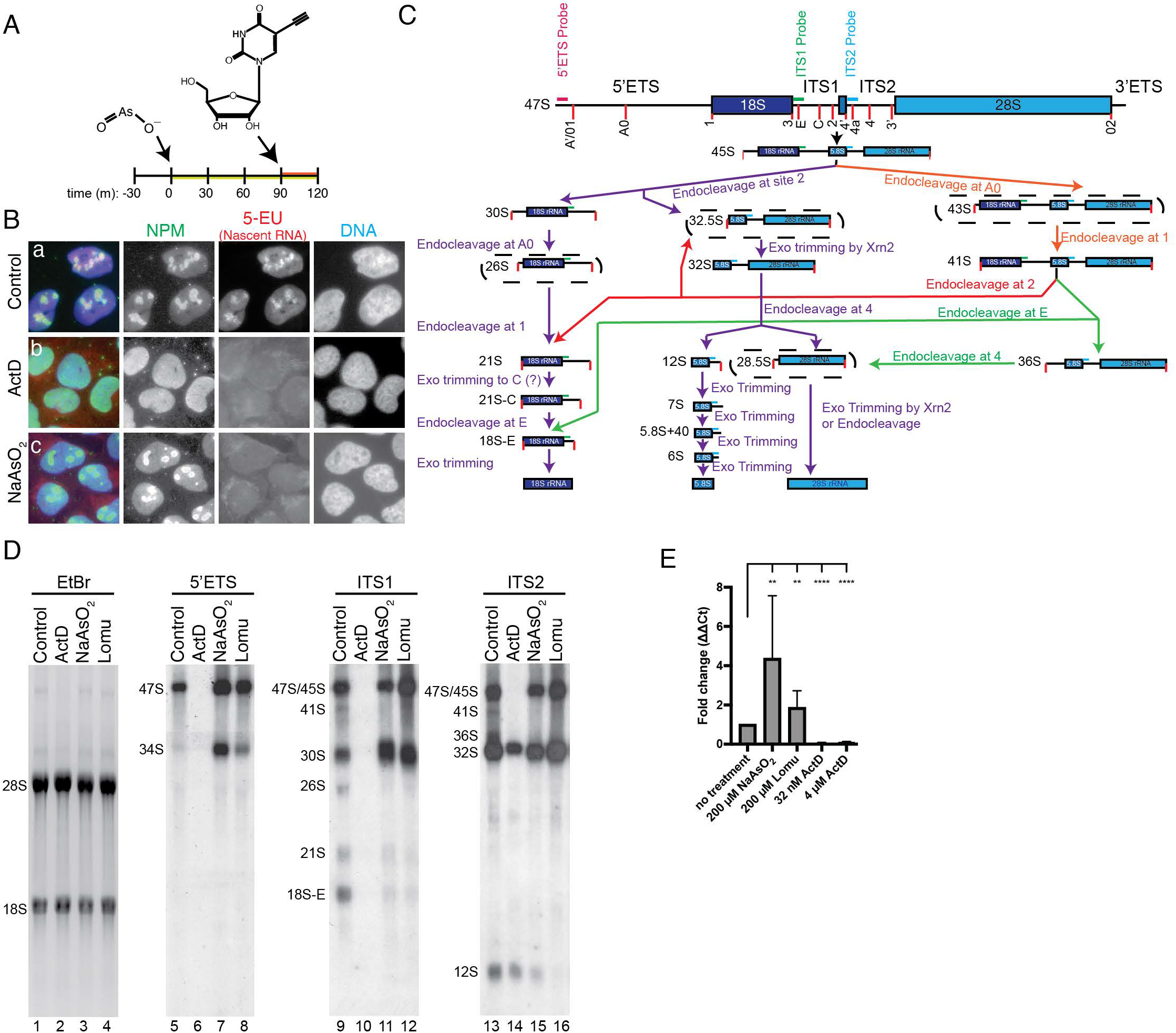
rRNA processing is inhibited in response to ISR-activating stress. **(A)** Schematic of 5-EU metabolic labeling experiment. Cells were stressed for 1.5 hrs and then 5-EU was added for 30 minutes to monitor transcriptional output. **(B)** Transcription in the nucleolus has been inhibited after 2 hours of NaAsO_2_-induced stress as indicated by loss of 5-EU signal in the nucleolus. **(C)** Schematic of the maturation pathway of human rRNAs. Approximate locations of northern blotting probes are noted in magenta, green and blue. **(D)** Northern blotting of cellular RNA after indicated treatments. 5 μg of RNA was resolved on 1.2% HEPES-EDTA agarose gel, transferred to nylon membrane and probed with indicated northern blotting probes. NaAsO_2_ and lomustine result in accumulation of 47S pre-rRNA, the generation of the aberrant 34S product and the decrease in downstream precursors (18S-E, 21S, 26S, 41S, 12S). **(E)** qRT-PCR confirms increase in 47S pre-rRNA precursor after NaAsO_2_ and Lomustine treatment and the decrease in 47S levels after ActD treatment. Data analyzed by student t-test (** = p<0.01, **** p<0.0001).

The transcription factor p53 plays a major role in the maintenance of the nucleolus (Rubbi and Milner 2003; Woods et al. 2015). U2OS cells are a p53 positive cell line, so, to address the possibility that regulation of rRNA processing was dependent upon p53, we performed the same experiments in HeLa cells, which are p53-null due to overexpression of the Human Papillomavirus E6 gene (Liu et al. 1999). We found that regulation of processing was independent of p53 status as it occurs in both the p53-positive U2OS cells and the p53-deficient HeLa cells. We also wanted to address whether or not this regulation was specific to cancer cells, so we assayed non-cancerous human RPE-1 cells that had been immortalized with hTERT. Again, we found the regulation as in the cancerous U2OS and HeLa cells (**Figure S1C**). Finally, the A’/01 processing site is conserved in mice, so we wanted to assay if the regulation of processing was also conserved across evolution. Using NIH3T3 cells, we found the same increase in 47S rRNA, loss of downstream precursors and generation of 34S pre-rRNA intermediate (**Figure S1D**). Finally, we show that loss of downstream precursors and generation of the 34S fragment occurs in a dose-dependent and time-dependent manner (**Figure S1E**). Therefore, stress-dependent regulation of rRNA processing is an evolutionarily conserved mode of stress response that occurs in response to various stresses, in a p53-independent manner in both transformed and untransformed cells.

The rate and efficiency of RNA processing strongly affect the rate of transcription and vice-a-versa (Schneider et al. 2007). That is, inhibition of rRNA processing can feedback to reduce rRNA transcription. The accumulation of 47S rRNA coupled with the loss of transcription after 90 minutes of stress suggested that rRNA processing was inhibited, resulting in accumulation of the primary transcript (**Figure 1 D – E**), which eventually results in cessation of RNA transcription (**Figure 1A – 1B**). Thus, we complemented our analysis of processing by ^32^P-metabolic labeling (**Figure 2A - C**). Here, cells were stressed and [^32^P]-*ortho*-phosphoric acid was added to monitor rRNA transcription and processing before chasing with cold, NaAsO_2_-containing media for 2.5 hours (**Figure 2A**) in accordance with published protocols (Pestov et al. 2008). In control conditions, initial precursors (47S, 45S), intermediates (30, 32S) and mature rRNAs (28S, 18S) were all observed and matured over the 3.5-hour time-course (**Figure 2B**, ln 15 - 21). However, in agreement with northern blotting data, after NaAsO_2_ treatment, initial precursors were generated, but these never matured to 18S and 28S rRNAs (**Figure 2B**, ln 22 - 28). We did observe RNA species approximately the size of the 30S and 32S precursors that our northern blotting data, particularly our dose-response data (**Figure S1E**), would suggest that it is the 34S fragment, and not a canonical processing intermediate. However, this assay does not allow us to identify this fragment unambiguously. Regardless, these data confirm that rRNA processing is inhibited by stress rather than directly targeting transcription. Treatment with the transcriptional inhibitor, Actinomycin D, completely abolished new synthesis of rRNA (**Figure 2C**). We propose that inhibition of pre-rRNA processing serves to “pause” ribosome biogenesis during a stress response. This occurs coordinately with inhibition of ribosomal protein biogenesis caused by eIF2α phosphorylation. However, for this “pausing” to be an effective stress response strategy, the accumulated 47S rRNA should be utilized when stress has passed. Upon NaAsO_2_ washout, eIF2α is rapidly dephosphorylated and translation resumes (Novoa et al. 2003). To assay if this occurs, we performed the same ^32^P-metabolic labeling experiments, but followed the labeled RNA after removing NaAsO_2_. The timing of this experiment is critical for accurate interpretation (**Figure 2E**). Cells were stressed with NaAsO_2_ concurrently with ^32^P labeling as before. ^32^P was removed and excess cold phosphate was added. Finally, NaAsO_2_ was removed and we monitored rRNA biogenesis over a 9-hour time course. This experimental setup demonstrated that rRNA was matured under control conditions as 18S and 28S rRNAs were generated by 2 hours. This process was stalled by NaAsO_2_ as demonstrated previously, but after washout, the radiolabeled pre-rRNA began to accumulate as mature rRNAs (**Figure 2E**). The final amount of pre-rRNA that matured to 18S and 28S was significantly lower than under control conditions because of the stalling in rRNA synthesis during stress. Therefore, during a stress response, rRNA processing stalls upon inhibition of processing at the A’/01 processing site, leading to the accumulation of 47S pre-rRNA. When stress resolves, this stored 47S pre-rRNA re-enters the biogenesis pathway leading to the production of mature rRNAs.

**Figure 2.**
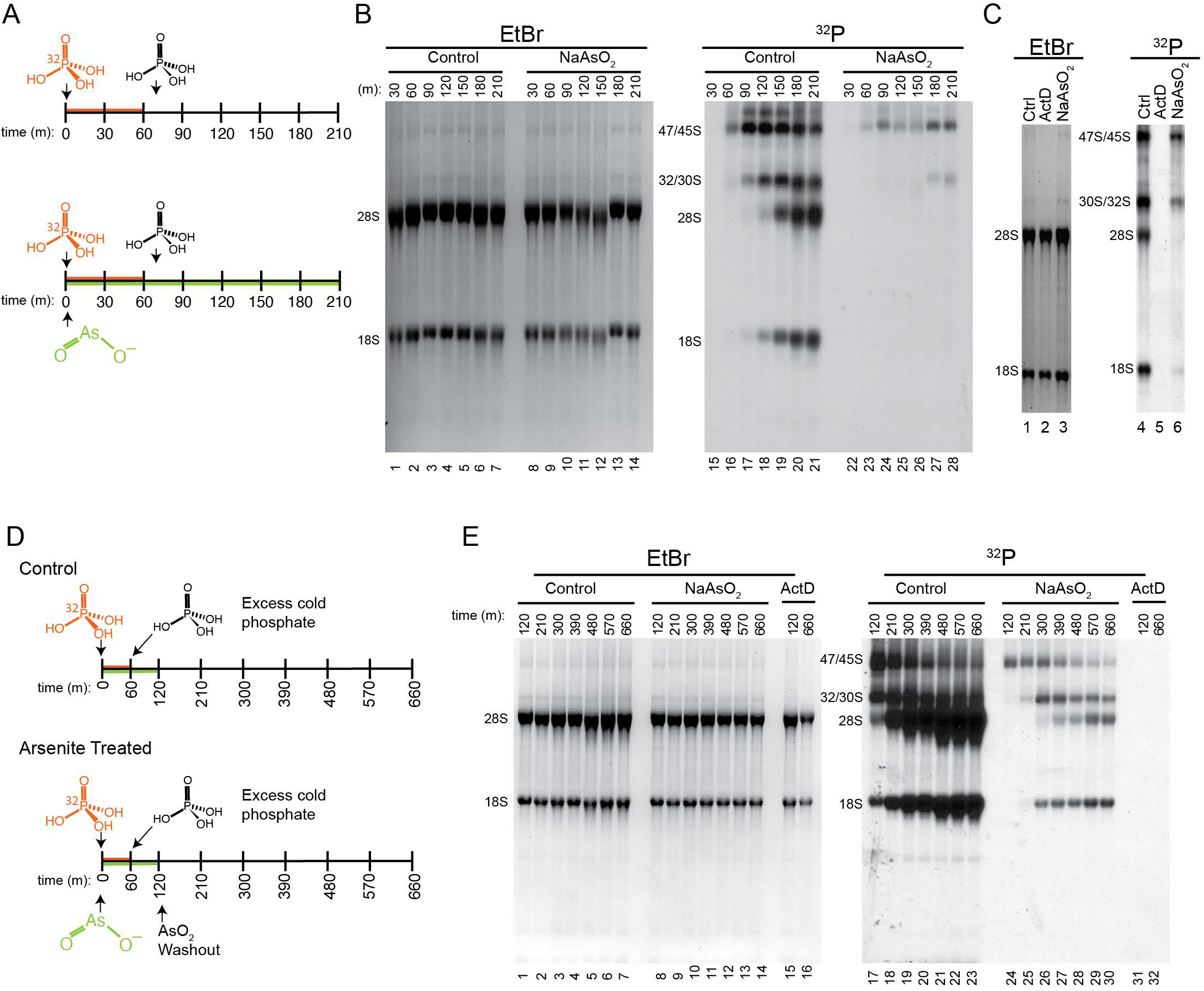
Stalled rRNA processing intermediates are utilized following a return to homeostasis. **(A)** Schematic of [^32^P]-*ortho-*phosphoric acid labeling experiment. Cells were stressed for 10 minutes before addition of [^32^P] to monitor rRNA transcription **(B)** NaAsO_2_-treatment results in inhibition of rRNA processing. Large precursor rRNAs are generated, but the never mature into 18S and 28S rRNAs. **(C)** End-point metabolic labeling with cells treated with NaAsO_2_ or Actinomycin D (ActD). Whereas NaAsO_2_ inhibits rRNA processing, ActD prevents rRNA transcription. **(D)** Schematic of [^32^P]-*ortho-*phosphoric acid labeling recovery experiment. Cells were stressed and labeled as in (A), but following 2 hours, NaAsO_2_ was washed out and cells were allowed to recover over the indicated time course. **(E)** After NaAsO_2_ washout, rRNA that was labeled during the initial phases of stress response re-entered the rRNA maturation pathway resulting in mature 18S and 28S rRNAs. Treatment with ActD prevents labeling of rRNA at the beginning of the time-course (120 m) or after washout (660 m).

The transcription and processing of rRNAs and the maturation of ribosomes occur in the nucleolus. Since this pathway is regulated in response to stress, we sought to investigate further if stress triggered any changes to the nucleolus. While untransformed cells typically have 2 – 3 nucleoli, it has been long established that cancer cells often exhibit an increased number of nucleoli that are often of increased size, an observation that has been shown to be correlated with proliferative potential (Montanaro et al. 2008). Indeed, human osteosarcoma U2OS cells examined in this study have between 3 and 5 nucleoli on average as monitored by immunostaining against NPM (**Figure 3Aa & 3B**). Treatment of these cells with ActD, significantly reduces the size or abolished the nucleoli as monitored by NPM (**Figure 3Ab & 3C**) without affecting the nucleolar number of cells that retained nucleoli on a per-cell basis (**Figure 3B**). RNA and RNA-binding proteins (RBPs) are major drivers of LLPS in non-membranous organelles (Lin et al. 2015; Protter et al. 2018; Van Treeck et al. 2018). Thus, as transcription inhibition reduces RNA levels in the nucleolus, nucleolar structure is disrupted. However, as previously shown, this is not coincident with activation of the ISR (**Figure S1A**), which is further exemplified here by lack of stress granule formation as monitored by eIF3B localization (**Figure 3A**). Upon treatment with NaAsO_2_ or lomustine we induced the formation of stress granules; however, NPM-positive nucleoli remained (**Figure 3Ac-d**). NPM is but one marker of nucleoli which mass spectrometry analysis has determined contains over 270 protein (Andersen et al. 2002). These proteins are distributed throughout three subcompartments within the nucleoli: Granular component (GC), Dense Fibrillar component (DFC) and Fibrillar component (FC). To ensure that NPM, a GC resident protein, was not unique in its nucleolar retention, we analyzed 11 additional nucleolar components by immunofluorescence representing members of each subcompartment and found no appreciable change in localization; however, future research will be needed to analyze more subtle changes in nucleolar substructures. (**Figure S2A – H**). As with regulation of rRNA processing, using HeLa cells, we also show that persistence of the nucleolus during stress is independent of p53 (**Figure S2J**).

**Figure 3.**
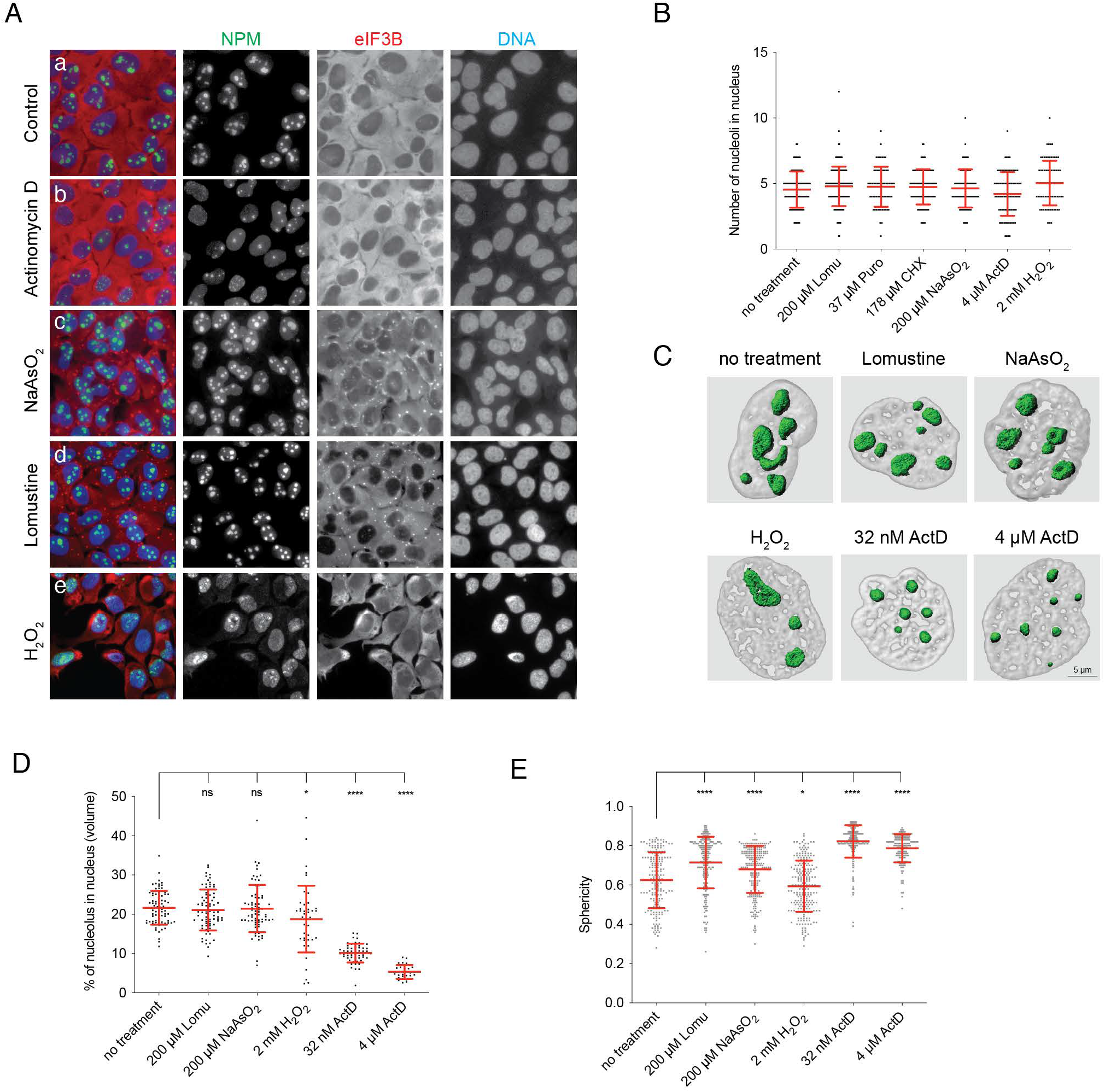
Nucleolar structure is maintained upon activation of the Integrated Stress Response. **(A)** Immunofluorescence of untreated U2OS osteosarcoma cells **(Aa)** or treated with Actinomycin D **(Ab)**, NaAsO_2_ **(Ac)**, Lomustine H_2_O_2_ **(Ad)**, or H_2_O_2_ **(Ae)**. Nucleolar integrity was monitored by staining for nucleophosmin (NPM, green) and activation of the ISR was monitored by analysis of stress granule formation using eIF3B (red). **(B)** Nucleolar number is unaffected by stress response on a per cell basis. Cells were treated with indicated stresses and the number of retained nucleoli per nuclei were manually counted using NPM as a nucleolar marker. **(C)** Representative images of Imaris 3D reconstruction after indicated treatments. **(D)** Nucleolar volume is unaffected after ISR-activating stresses (NaAsO_2_ and Lomustine), while volume is significantly decreased after mTOR-inactivating H_2_O_2_ treatment and ActD, which inhibits transcription. Data analyzed by student t-test (* = p<0.05, **** p<0.0001). **(E)** Sphericity of nucleoli was determined following 3D reconstruction. ISR-activating stresses results in a more spherical nucleolar structure, while mTOR-inactivating H_2_O_2_ treatments results in a statistically significant decrease in sphericity Data analyzed by student t-test (* = p<0.05, **** p<0.0001).

It is worth noting that these data seemingly contradict a previous report that showed that oxidative stress initiated by H_2_O_2_ inhibits rRNA synthesis via phosphorylation of TIF-1A, a basal polymerase I transcription factor (Mayer et al. 2005). Thus, we sought to address this apparent discrepancy. In doing so, we repeated previously reported data by showing that H_2_O_2_-induced oxidative stress disrupts nucleolar architecture (**Figure 3Ae, 3B - D**). However, our previous data showed that H_2_O_2_ not only activates the ISR, but also potently inhibits mTOR (Emara et al. 2012) as indicated by an increase in non-phosphorylated 4EBP1 (**Figure S1A)**. Since mTOR is required for TIF-1A activation, the different targets of H_2_O_2_- and NaAsO_2_-induced oxidative stress explains the apparent contradiction (Mayer et al. 2004). We conclude that persistence of the nucleolus during stress is a feature of ISR-activating oxidative stress when mTOR is active, but not when it is inactive.

To further analyze nucleolar size and morphology, we employed Imaris imaging software to reconstruct 3D models of nucleolar structure under various cellular conditions to make parametrical analysis of the organelles and, in consequence, more precise measurements (**Figure 3C**). This analysis revealed that upon the initiation of a stress response, there was no apparent change in nucleolar volume, in contrast to inhibition of transcription by ActD (**Figure 3D**). Immunofluorescence also suggested a change in nucleolar morphology upon induction of a stress response, namely adoption of more rounded morphology. We specifically quantified this by measuring sphericity as a ratio between the radius of an inscribing and circumscribing circle of the nucleoli. Thus, the more spherical the nucleoli, the closer the sphericity will be to 1, and the more oblong, the closer the shape will be to 0. Upon analysis, we showed a statistically significant increase in sphericity after NaAsO_2_ and lomustine treatment, suggesting a change in the biophysical dynamics of this organelle (**Figure 3E**).

Since RNA is a contributing factor of nucleolar assembly via an RNA-driven liquid-liquid phase separation, we next sought to determine if this unprocessed rRNA was retained within the nucleolus, as would be suggested by the persistence of nucleoli and the utilization of stored pre-rRNA after stress (**Figure 2 & 3**). We performed RNA FISH in conjunction with immunofluorescence to analyze the localization of rRNA using Imaris 3D reconstruction. Following stress, 47S rRNA was retained within the nucleolus which aids in the preservation of nucleolar structure (**Figure 4A**). Maintenance of this RNA in the nucleolus (**Figure 4 A**), inhibition of processing (**Figure 1 - 2**) and increased nucleolar sphericity (**Figure 3E**) would suggest that the unprocessed RNA is being stored in the nucleolus thereby effecting nucleolar dynamics during stress. To assess this, we performed fluorescence recovery after photobleaching (FRAP) of nucleoli under stressed and unstressed conditions. Cell lines stably expressing mCherry-Nol9, an rRNA processing factor, GFP-RPL7A, a ribosomal protein, and mCherry-NPM were generated (**Figure S3A**). We performed FRAP on control cells or cells treated with NaAsO_2_ for 2 hours (**Figure 4B – E, Figure S3B – C)**. Since the nucleolus is an extremely active organelle, we were not surprised to find that fluorescence recovered nearly completely 45 seconds post-bleaching for each of the three tagged proteins, with NPM recovering after only 10 – 12 seconds post-bleach. However, NaAsO_2_ treatment severely diminished nucleolar dynamics. We observed almost no recovery over the same timescale. Particularly striking was the change in recovery of Nol9, an rRNA processing factor. These data confirm that in response to stress, the nucleoli serve as storage sites for unprocessed rRNA. The presence of this rRNA aids in the persistence of nucleoli.

**Figure 4.**
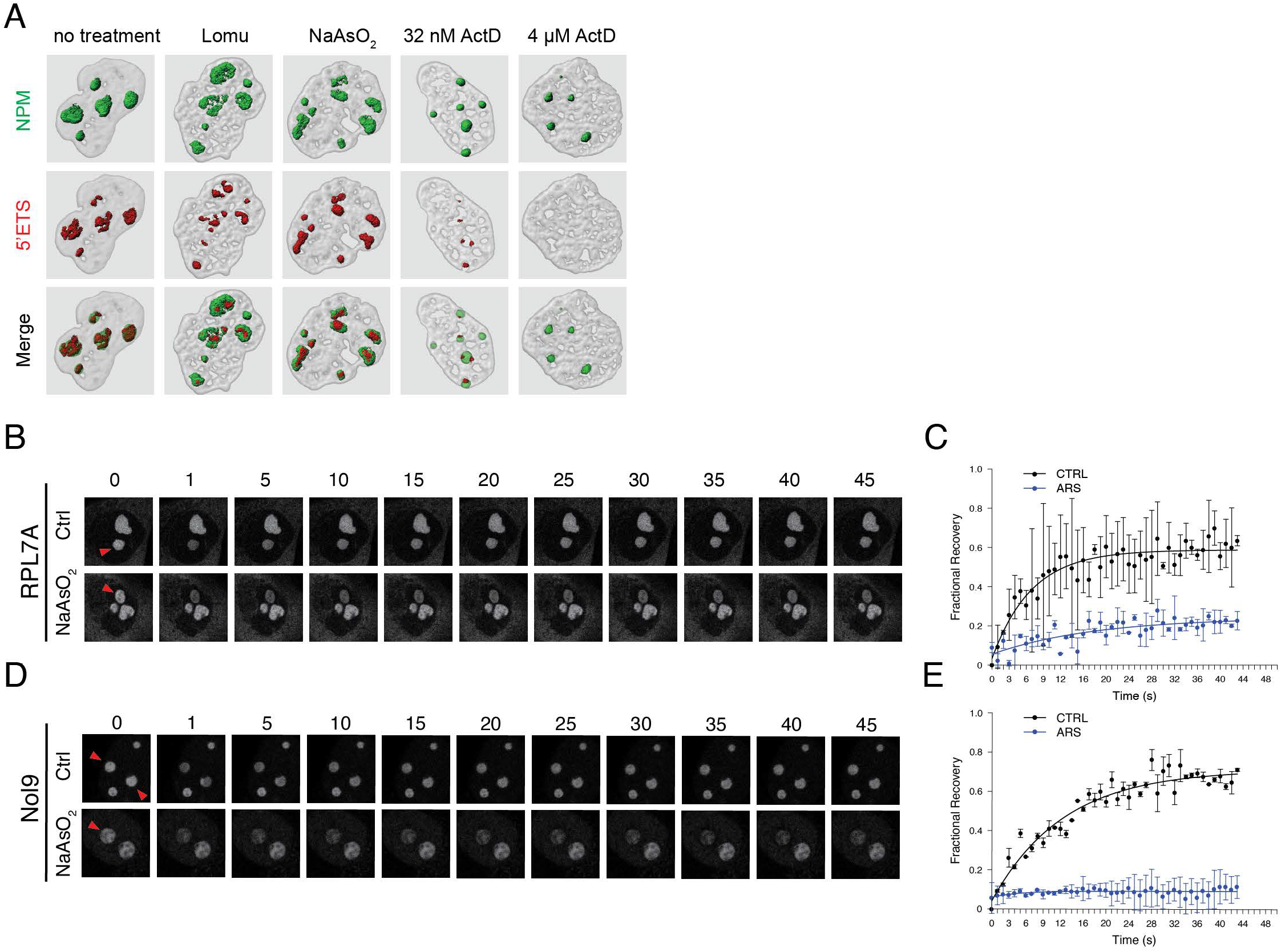
Unprocessed rRNA is stored in the nucleolus resulting in perturbed nucleolar dynamics. **(A)** 3D reconstructions of nucleoli after indicated treatment using NPM and FISH probes to 5’ETS. Results demonstrate that unprocessed rRNA is retained within the nucleolus during a stress response. **(B – E)** Fluorescence recovery after photobleaching of RPL7A **(C – C)** or Nol9 **(D – E**) under basal and stressed conditions. Under nominal conditions, the bleached fluorescence signal rapidly recovers as expected by an active organelle. However, fluorescence recovery is severely perturbed during a stress response, consistent with the inhibition of rRNA processing. Red arrowheads denote the photobleached nucleoli.

We next sought to determine if regulation of rRNA processing in response to stress is a component of the ISR. To begin, we analyzed transcription two hours post-stress using 5-EU as before (**Figure 1A**). Formation of stress granules is typically seen as a proxy for ISR activation (Kedersha et al. 2013). Thus, we treated cells with levels of NaAsO_2_ below the level that fully induces SG formation (**Figure 5A**). At 75 μM NaAsO_2_, approximately 50% of cells have visible SGs as monitored by eIF3B staining. However, regardless of whether or not a cell has SGs or not, nucleolar transcription has ceased, suggesting that regulation of rRNA is independent of the ISR. To further explore whether activation of ISR is a requirement of stress-dependent rRNA regulation, we knocked out heme-regulated inhibitor kinase (HRI/EIF2AK1) using CRISPR/Cas9 (**Figure S4**). We have previously shown that CRISPR/Cas9 knockout of HRI renders cells unresponsive to NaAsO_2_ with regards to eIF2α phosphorylation, translational repression and SG formation (Aulas et al. 2017). Therefore, we sought to determine whether rRNA processing was still regulated if the ISR was inhibited by deletion of HRI. SGs form in wild-type U2OS cells in response to NaAsO_2,_ while ΔHRI U2OS cells fail to form SGs (**Figure 5B**). However, 5-EU labeling of nascent RNA reveals transcriptional shutoff of nucleolar RNA synthesis two hours post-stress in both WT and ΔHRI cells. To further explore the connection, or lack thereof, between the ISR and rRNA regulation, we processed WT and ΔHRI cells for northern blotting analysis before and after stress. We still found the generation of the 34S fragment after NaAsO_2_ treatment (**Figure 5C and D**) and a decrease in downstream rRNA precursors, indicating that inhibition of pre-rRNA processing occurs in a stress-dependent manner, independent of the ISR. Finally, we completed ^32^P pulse-chase experiments to monitor rRNA processing. We found that, as in WT U2OS cells, 47S rRNA was generated in ΔHRI U2OS cells, but this never matured to 28S or 18S rRNA after induction of a stress response (**Figure 5D**). Therefore, regulation of rRNA biogenesis in response to stress is a parallel, but independent, pathway to the ISR. We term this previously undescribed pathway the “Ribosome Biogenesis Stress Response (RiBiSR).”

**Figure 5.**
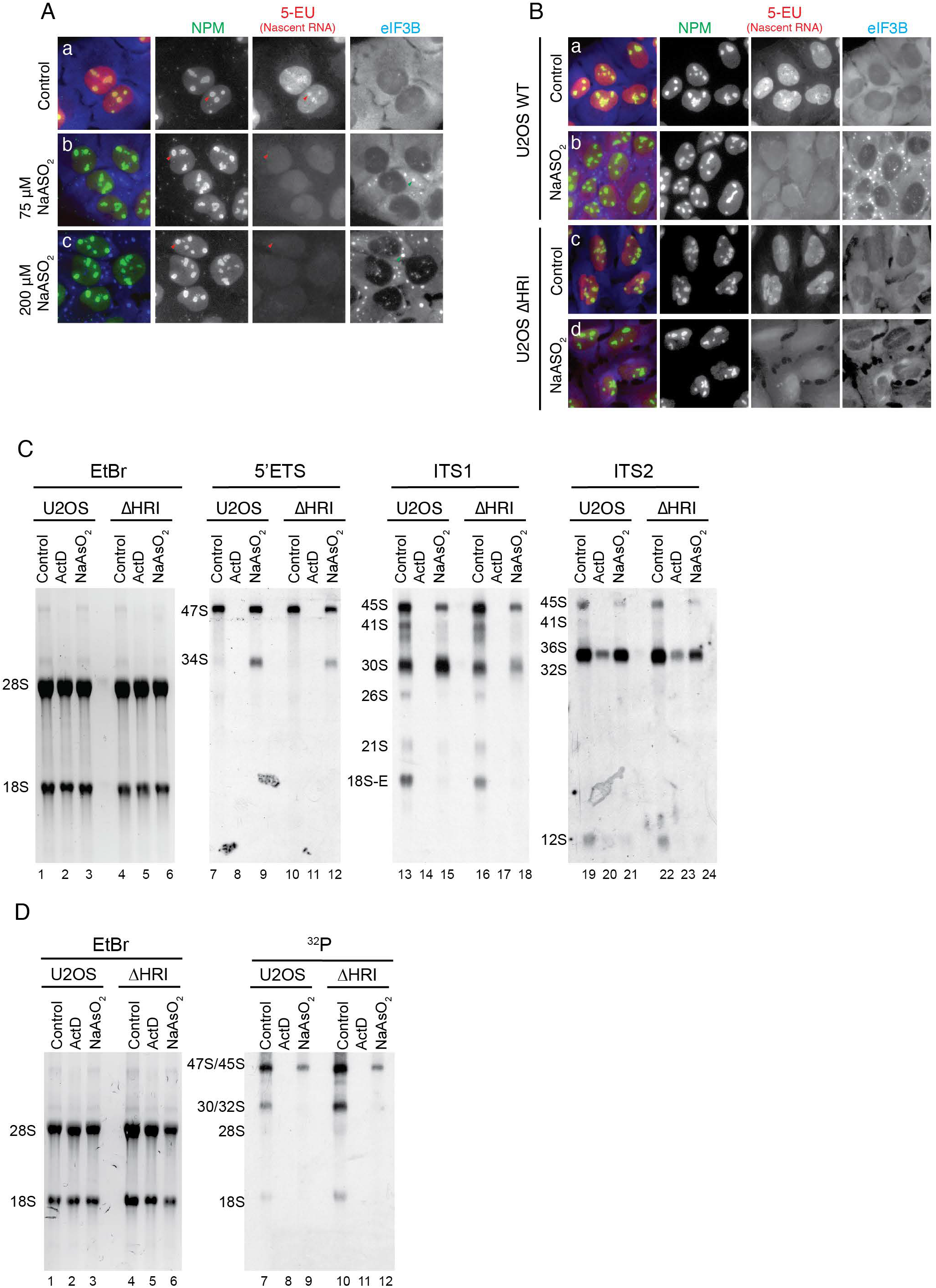
Regulation of rRNA processing in response to stress is parallel to, but not dependent upon, the ISR. **(A)** 5-EU metabolic labeling was done as before with 0 μM **(Aa)**, 75 μM **(Ab)** or 200 μM NaAsO_2_ **(Ac)**. Formation of stress granules, as monitored by eIF3B, was used as a proxy for activation of the ISR. rRNA transcription was impaired after 2 hours of stress regardless of the formation of stress granules. Red arrowheads denote the nucleoli and green arrowheads denote stress granules. **(B)** 5-EU metabolic of **WT (Ba-b)** or ΔHRI **(Bc-d)** cells untreated or treated with NaAsO_2_. ΔHRI cells fail to form stress granules as translation is not regulated in these cells in response to stress. However, rRNA transcription is still inhibited 2 hours post-stress induction. **(C)** Northern blotting of rRNA in WT and ΔHRI U2OS cells. Both cells respond to NaAsO_2_ by generating the 34S fragment, diagnostic of processing inhibition (ln 9 & 12) and leading to the reduction of downstream precursors (ln 15, 18, 21, & 24). **(D)** [^32^P]-metabolic labeling of WT and ΔHRI cells demonstrate an inhibition in rRNA processing in response to cell stress despite a failure to inhibit translation.

We have argued that RiBiSR serves to maintain cellular homeostasis during stress by preventing unnecessary production of rRNA when ribosomal protein synthesis has been inhibited in the cytoplasm. Further, we argue that it functions to maintain the balance between ribosomal protein synthesis and rRNA synthesis, thereby preserving nucleolar integrity and protecting against nucleolar stress. To test this hypothesis and understand what the consequences of disrupting this balance are, we treated cells with puromycin (Puro) or cycloheximide (CHX). Both Puro and CHX are pharmacologic inhibitors of translation that do not activate the ISR or inactivate mTOR (**Figure S1A**). They both directly target the translational machinery: Puro triggers premature translation termination while CHX stalls ribosome elongation. Therefore, treatment with these drugs would inhibit *de novo* ribosomal protein synthesis without directly targeting rRNA synthesis. Upon treatment of cells with CHX and Puro, despite inhibiting translation through different mechanisms, we found similar results. We began by analyzing nucleolar dynamics by FRAP (**Figure 6A – D, Figure S5A – B**). We argued that stress-induced decline in nucleolar dynamics served to maintain nucleolar structure after the inhibition of ribosomal protein synthesis. However, upon translation inhibition by Puro or CHX, we found no significant change in nucleolar dynamics irrespective of the fact that new ribosomal proteins are not being delivered to the nucleolus (judging by either RPL7A (**Figure 6A – B**) or Nol9 (**Figure 6C – D**)). Next, we analyzed rRNA by northern blotting and found that 47S rRNA was still present without the production of the stress-dependent 34S fragment (**Figure 6E**). However, there was a reduction in the initial 47S rRNA precursor (**Figure 6E**, ln 9 & 10). Additionally, downstream precursors (e.g., 12S, 21S, 18S-E) are reduced upon treatment with Puromycin and CHX, consistent with the reduction of 47S, but not abolished as it is seen with NaAsO_2_. This is consistent with cells maintaining the balance between ribosomal protein and rRNA synthesis. However, pharmacological inhibition of ribosomal protein synthesis results in an inhibition of 47S synthesis, not processing. Finally, we have argued that stress-dependent regulation of rRNA processing preserves nucleolar integrity when ribosomal protein synthesis has been inhibited. As CHX and puromycin do not induce regulated inhibition of rRNA processing, we sought to determine the effect of continued rRNA production in the absence of ribosomal protein synthesis on nucleolar morphology by immunofluorescence (**Figure 6F**). As we found previously, nucleolar structure was preserved after NaAsO_2_-induced stress, with individual nucleoli adopting a more spherical morphology, concurrent with the formation of stress granules in the cytoplasm (**Figure 6Fb**). In contrast, neither CHX nor puro induces SG formation, but there was a dramatic effect on nucleolar morphology (**Figure 6Fc-d**). Intact nucleoli adopted a “ragged” morphology. However, more striking, was that the nucleoli in a subset of cells became fragmented indicating a total loss of nucleolar integrity. The fact that there was no compensatory nucleolar response to the loss of ribosomal protein production likely contributes to the resultant nucleolar fragmentation. These results demonstrate the importance of RiBiSR for maintenance of cellular homeostasis under stress.

**Figure 6.**
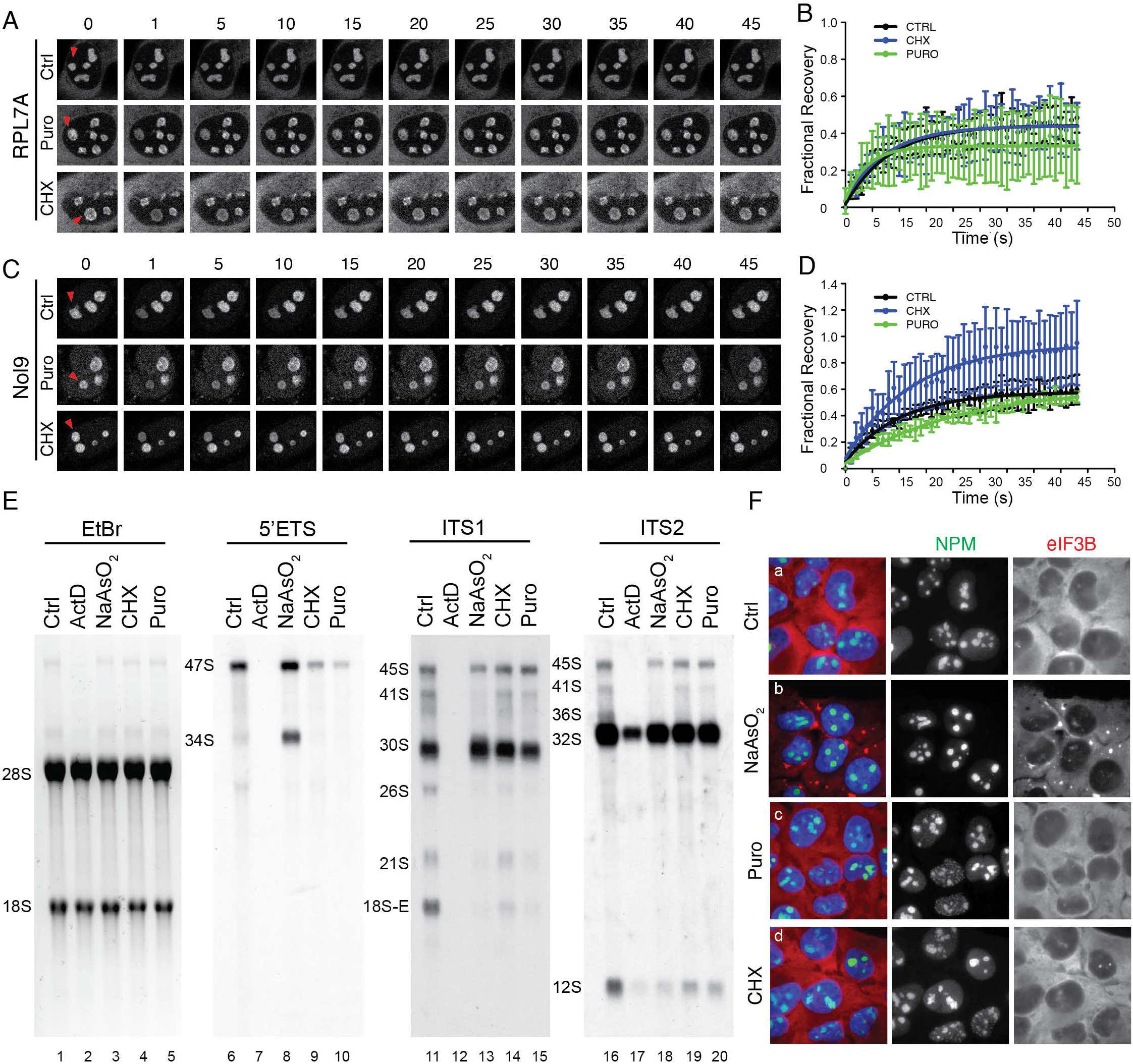
Failure to regulate rRNA processing with translation inhibition results in nucleolar fragmentation. **(A - D)** FRAP of U2OS cells stably expressing RPL7A **(A – B)** or Nol9 **(C – D)** untreated or treated with cycloheximide or puromycin to repress translation without inducing a stress response. Despite inhibition of translation, nucleolar dynamics remain unaltered indicating that previously identified alteration in nucleolar dynamics is not a result of translation inhibition. Red arrowheads denote photobleached nucleoli. **(E)** Northern blotting of rRNA after treatment with CHX and puro demonstrates that 47S rRNA and 34S fragment do not accumulate. Instead, there is a reduction in 47S levels (ln 9 & 10) commensurate with the inhibition of ribosomal protein synthesis. **(F)** After indicated treatments, nucleoli of U2OS cells were analyzed by immunofluorescence by staining for NPM. Stress granule formation was monitored by eIF3B localization. As shown previously, NaAsO_2_ results in stress granule formation and maintenance of nucleolar structure. However, translational inhibitors that do not trigger a stress response (CHX and Puro) result in fragmentation of nucleoli.

## DISCUSSION

Our work reveals a novel stress-response pathway in mammalian cells that we have termed RiBiSR. This pathway is independent of the ISR, which functions in the cytoplasm to initially inhibit translation initiation. This novel program regulates the biogenesis of ribosomes, amongst the most energy-intensive, pro-growth processes in the cell. The goal of any stress response is to promote survival until the return to homeostasis. This is largely accomplished by the reallocation of energy reserves away from pro-growth activities towards pro-survival activities. Since the majority of cellular energy is devoted to ribosome biogenesis (Warner et al. 2001), it is unsurprising that this pathway is under substantial regulatory pressure during a stress response. Much work has gone into demonstrating how protein synthesis and specifically the translation of ribosomal proteins is regulated in response to stress (Ivanov et al. 2011; Meyuhas and Kahan 2015). Others have previously shown that inactivation of mTOR in response to nutrient starvation, downregulates the transcriptional rate of rRNA genes (Mayer et al. 2004). Additionally, in yeast, TORC1 signaling has also been implicating in regulating the processing of rRNA in response to certain stresses (Kos-Braun et al. 2017). Here, we uncover a novel mechanism that is responsive to acute stress that functions via regulation of rRNA processing. Our data show that upon activation of the ISR, the nucleolus is maintained by inhibiting rRNA processing rather than inhibiting rRNA transcription. The initial processing event, conversion of the 47S to 45S rRNA, is under tight stress-dependent regulation. Unlike other stress response pathways that regulate ribosome biogenesis, this pathway is not dependent upon mTOR, nor is it dependent upon ISR-mediated translation repression.

Importantly, this mechanism of regulation results in the preservation of the nucleolus. A major contributing factor to nucleolar structure is driven by a liquid-liquid phase separation (LLPS). FBL, the 2’O methyltransferase, aids in the establishment of an LLPS by binding nascent rRNAs and sorting them into nucleolar compartments (Yao et al. 2019). Additionally, the multimerization of NPM plays a critical role in establishing nucleolar structure (Mitrea et al. 2018). Since RNA is a major driver of LLPS [Reviewed in (Van Treeck and Parker 2018)], stresses or cellular conditions that inhibit rRNA transcription result in the dissolution of nucleolar structure due to the absence of rRNA in this compartment (ActD, **Figure 3**).

Nuclear bodies, including the nucleolus, are often formed around regions of high transcriptional output and serve to concentrate factors involved in RNA metabolism (Reviewed in (Dundr and Misteli 2010)). In addition to the nucleolus, this has been shown for other nuclear bodies, including the histone locus body (Tatomer et al. 2016) and Cajal bodies (Xu et al. 2005). Therefore, if a stress response were to result in the destruction of the nucleolus, this would only further hamper cell viability as the cell recovers from a cellular insult. Nucleolar components would be dispersed throughout the nucleoplasm and need to be relocalized to begin ribosome biogenesis. We propose that by maintaining nucleolar structure, an efficient return to homeostatic conditions is ensured upon the cessation of cellular insult without wasting an additional energy in reassembling this nuclear body.

We propose that this mechanism has parallels to the role of translation inhibition following activation of the ISR that results in phosphorylation of eIF2α by one of four stress-responsive kinases. This pathway culminates in the stalling of translation at the initiation stage, but, importantly, this does not result in the degradation of mRNAs or the disassembly of translation initiation complexes. Instead, stalled 48S pre-initiation complexes and mRNAs are stored in SGs, which themselves are liquid-liquid phase condensates. As the stressed state resolves, stored mRNAs can re-enter the pool of translating mRNAs. Stress-dependent regulation of rRNA processing does not destroy rRNA, but stalls its biogenesis at the initial stages, similar to inhibition of translation initiation. Additionally, regulation of processing preserves the localization of ribosome biogenesis machinery within the nucleolus. This is in contrast to mTOR-inactivating stresses that affect both rRNA transcription and mRNA translation. Starvation conditions are particularly potent inhibitors of mTOR. With regards to translation, this results in the dephosphorylation of 4EBP1 proteins which disrupt the eIF4F and pre-initiation complex formation. Similarly, mTOR inactivation results in inhibition of transcription of rDNA genes through regulation of TIF-1A/RRN3 (Mayer et al. 2004). Here, this results in disassembly of the nucleolus which disperses rRNA processing enzymes throughout the nucleoplasm. In yeast, TORC1 signaling has been shown to affect the processing of the 35S rRNA by causing a switch in the site of ITS1 processing (Kos-Braun et al. 2017).

Our data also suggests that conversion of the 47S to 45S rRNA is an obligatory step in rRNA biogenesis. When the pre-RNA matures to the 45S intermediate, there are multiple pathways through which the RNA intermediate can mature (reviewed in (Mullineux and Lafontaine 2012)). However, our results demonstrate that, when stalled as the 47S rRNA, there is no alternative pathway that leads to mature 18S, 5.8S and 28S rRNAs. The fraction of 47S that gets processed to the 34S is not matured despite the fact that this represents the same RNA species as 30S rRNA except for the 5’ extension (i.e. 1 – 01 fragment). It is tempting to speculate that this region contains an inhibitory sequence that prevents further processing. Alternatively, all rRNA processing enzymes could be simultaneously inhibited, but this would necessitate a here-to-fore unforeseen level of rRNA processing regulation. Another outstanding question is whether the 34S rRNA has any biological function. It is generated in a dose- and time-dependent manner during a stress response (**Figure S1E**). This rRNA species has been observed upon the knockdown of rRNA processing components necessary for small subunit biogenesis (e.g. FBL) (Tafforeau et al. 2013). Our FISH data demonstrates that this fragment is retained in the nucleolus, but whether it plays a functional or structural role there is unknown.

Importantly, the enzyme responsible for cleavage at the A’/01 site in the 5’ETS is unknown. This initial processing site is conserved in mice and humans (Mullineux and Lafontaine 2012), as is the stress-dependent regulation (**Figure S1**). Processing here, and at the 02 site at the 3’ end of the pre-rRNA, convert the 47S rRNA to the 45S rRNA. The identity of this enzyme (or enzymes) and how its activity is regulated in response to stress are critical to gain a full understanding of the cellular stress response. We show that this pathway is parallel to, but not dependent upon ISR activation (**Figure 5**). This suggests the existence of an alternative stress-responsive pathway found in the nucleus or nucleolus. However, the identity of the players in this pathway remain to be uncovered.

## MATERIALS AND METHODS

### Antibodies

TIAR (Santa Cruz, sc-1749), NPM (Santa Cruz, sc-70392), FBL (Cell signaling Technology, 2639), RPA194 (Santa Cruz, sc-4669), eIF2α (Santa Cruz, sc-133132), phospho-eIF2α (AbCam, 131505), 4EBP (Cell Signaling Technology, 9454), non-phospho-4EBP (Cell Signaling Technology, 4923S), TIF-1A/RRN3 (Santa Cruz, 2c-390464), RPL7A (Cell Signaling Technology, 2415) Nol9 (Protein Tech Group, 16083-1-AP), UBF (Santa Cruz, sc-13125), eIF3B (Santa Cruz, sc-16377)

### Cell culture and drug treatment

U2OS and HeLa cells were maintained in DMEM supplemented with 10% Fetal Bovine Serum and Penicillin/Streptomycin. NIH3T3 were maintained in DMEM supplemented with 10% Bovine Calf Serum and Penicillin/streptomycin in a humidified 37°C/5% CO_2_ incubator. NaAsO_2_ (Sigma), Lomustine (Selleckchem), H_2_O_2_ (Fisher), Actinomycin D (Arcos Organics) were added for indicated times at indicated concentration. Where not noted, NaAsO_2_ was treated at 200 μM.

### Epifluorescence Immunofluorescence

Cells were fixed and processed for fluorescence microscopy as described (Lyons et al. 2016). Briefly, cells were grown on glass coverslips, stressed as indicated and fixed with 4% paraformaldehyde in PBS for 15 minutes followed by 10 minutes post-fixation/permeabilization in −20°C methanol. Cells were blocked for 1 hour in 5% horse serum/PBS. Primary and secondary antibody incubations were performed in blocking buffer for 1 hour with rocking at room temperature. Secondary antibodies (Jackson Laboratories) were tagged with Cy2, Cy3 or Cy5. Following washes with PBS, cells were mounted in polyvinyl mounting media and viewed at room temperature using a Nikon Eclipse E800 microscope with a 40X Plan fluor (NA 0.75) or 100X Plan Apo objective lens (NA 1.4) and illuminated with a mercury lamp and standard filters for DAPI (UV-2A −360/40; 420/LP), Cy2 (FITC HQ 480/40; 535/50), Cy3 (Cy 3HQ 545/30; 610/75), and Cy5 (Cy 5 HQ 620/60; 700/75). Images were captured with SPOT Persuit digital camera (Diagnostic Instruments) with the manufacturers software and compiled using Adobe Photoshop 2020.

### Imaris 3D reconstruction and Parametric analysis

For Imaris imaging, cells were treated in the same way as for epifluorescence immunofluorescence. The crude Z-stack images were collected using Olympus FV10i confocal laser scanning microscope. Then, images were processed with Imaris 7.4.2 (Bitplane, UK) in order to get 3D reconstruction. The sizes of nuclei and nuclei were automatically calculated on the basis of Z-stacks composed of individual images. We then calculated the volume of all nucleoli in one nucleus in one cell. At least 100 cells from at least three independent experiments were taken for one analysis.

### Metabolic labeling

For 5-ethynyl uridine labeling experiments, experiments were conducted using Click-iT RNA Alexa Flor 488 Imaging Kit (ThermoFisher) according to manufacturer’s instructions. Cells were prepared as for immunofluorescence, but 30 minutes prior to fixation, 5-EU was added to a final concentration of 1 mM. For [^32^P]-metabolic labeling, cells were grown in a 6-well dish. Cells were starved for one hour in phosphate free DMEM containing 10% dialyzed FBS. Cells were then treated for 10 minutes with indicated compounds before addition of 20 μCi of ^32^P-*ortho*-phosphoric acid (Perkin Elmer). Cells were incubated for 1 hour with media containing drugs and ^32^P-*ortho*-phosphoric acid, before replacing with DMEM containing 10% FBS with indicated drug treatment for an additional 1.5 hours. RNA was harvested with Trizol reagent according to manufacturer’s instructions and resuspended in 30 μl of RNA loading dye. Ten microliters of RNA was loaded onto a 1.2% Agarose gel made with 1X H-E buffer (20 mM HEPES, 1 mM EDTA [pH 7.8] and 7% Formaldehyde. Gels were run overnight in 1X H-E buffer at 55 V with recirculation and then dried and exposed to film.

### Northern Blotting

Cells were grown to ∼ 80% confluency and RNA was extracted with Trizol reagent (Invitrogen) according to manufacturer’s instructions. 5 μg of extracted RNA was resuspended in RNA loading dye (7 μl of Formamide, 2 μl of Formaldehyde, 1 μL of 10X HEPES-EDTA Buffer, 1 μl of Ethidium bromide (0.4 mg/ml), 1 μl of bromophenol blue (0.5 mg/ml) and heated to 85°C for 10 minutes before placing on ice. Denatured RNA was loaded onto a 1.2% Agarose gel prepared in H-E buffer (20 mM HEPES, 1 mM ETDA [pH 7.8] and 7% formaldehyde. Gel was run in HEPES-EDTA buffer overnight at 55V with recirculation. The following day, the gel was subjected to mild alkaline treatment (10 minutes in 50 mM NaOH/10 mM NaCl), neutralization (10 minutes in 2.5X TBE) and equilibration in 2X SSC. RNA was transferred overnight by passive transfer to Hybond N_+_ Nylon membrane using 20X SSC. The following day, RNA was dried, crosslinked to membrane and pre-hybridized in 10 ml of UltraHyb pre-hybridization/hybridization solution (Invitrogen) for 1 hour at 60°C. Pre-hybridization solution was removed and 10 mL of fresh UltraHyb was added along with 15 μl of end labeled northern probe. Probe was incubated for 1 hour at 60°C and then overnight at 37°C. The following day, probes were washed twice with 2X SSC/0.1% SDS at 40°C and exposed to film. For stripping and re-probing, 50 mL of boiling 0.1X SSC/0.1% SDS was added to blots and allowed to come to room temperature twice.

### Preparation of northern blotting probes

Synthetic DNA oligonucleotides were prepared by Integrated DNA Technologies and resuspended to a final concentration of 6 μM in dH_2_O. For end-labeling, 1 μl of DNA was reacted with 2 μl of [^32^P]-γ-ATP (3000 Ci/mL) (Perkin-Elmer), 1 μl of 10X T4 PNK buffer, 1 μl of T4 PNK (NEB), and 14 μl of dH_2_O for 1 hour. The reaction was brought to 100 μl with dH_2_O and unincorporated nucleotides were removed by gel filtration through G-25 column (GE LifeSciences). The following oligonucleotide sequences were used for northern blotting: Human 5’ ETS (CGGAGGCCCAACCTCTCCGACGACAGGTCGCCAGAGGACAGCGTGTCAGC), Human ITS1 (GGCCTCGCCCTCCGGGCTCCGTTAATGAT), Human ITS2 (CTGCGAGGGAACCCCCAGCCGCGCA), Mouse 5’ETS (AAGCAGGAAGCGTGGCTCGGGGAGAGCTTCAGGCACCGCGACAGA), Mouse ITS1 (ACGCCGCCGCTCCTCCACAGTCTCCCGTTTAATGATCC), Mouse ITS2 (ACCCACCGCAGCGGGTGACGCGATTGATCG).

### qRT-PCR

For qRT-PCR cells were growing on 6-well plate. After treatment with corresponding drug, total RNA was isolated using Universal RNA/miRNA Purification Kit (EURx, Poland). RNA was quantified using Nanodrop ND-1000 (ThermoScientific, USA). cDNA was obtained using LunaScript™ RT SuperMix Kit (New England Biolabs, USA) and qPCR reactions were performed using Luna^®^ Universal qPCR Master Mix (New England Biolabs, USA) according to manufactural instructions. The qPCR reactions were done in CFX96 Touch Real-Time PCR Detection System (Bio-Rad, USA) and then analyzed in CFX Maestro Analysis Software (Bio-Rad, USA). The final graphs were prepared in GraphPad Prism 8. Primers for amplification spanned the A’/01 site ensuring that only 47S and not other precursors were amplified. The sequences were: GTGCGTGTCAGGCGTTC and GGGAGAGGAGCAGACGAG.

### FRAP

U2OS cells were transfected with plasmids containing tagged versions of RPL7A, Nol9 or NPM and selected with geneticin. Cells were grown in 4-compartment 35 mm glass bottom dish (Greiner) until ∼80% confluency. Cells were treated with indicated drugs for 2 hours before conducting FRAP on Zeiss as described previously (Kedersha et al. 2005). Acquired FRAP images were converted to parametric data with the use of ImageJ software and ImageJ macro programming language (Rueden et al. 2017). Initially, image stacks were subjected to drift correction using Manual Drift Correction Plugin implemented into the ImageJ macro source code (Manual drift correction (Fiji), Benoit Lombardot). Transformed sequence image stacks were aligned with ROI of each bleached area for subsequent parametric data acquisition. The output parametric data from each ROI was grouped into 3 categories, encompassing: bleached, background, and reference region. Results were exported to csv files and subsequently imported into R programming environment to facilitate calculations and plot generation (R Core Team (2019). R: A language and environment for statistical computing. R Foundation for Statistical Computing, Vienna, Austria (URL: https://www.R-project.org/). To eliminate noisy data Background Intensity Values (BG) were subtracted from Bleach Intensity Values (B) to obtain Bleach Corrected Values (B_corr) for each bleached region. Subsequently, Background Intensity Values were subtracted from Reference Intensity Values (Ref) to obtain Reference Corrected Values (Ref_corr). Final calculation was based on normalization of Bleach Corrected Values to Reference Corrected Values according to following equation: Normalized Bleach Corrected Values = Bleach Corrected Values/ Reference Corrected Values. Final data normalization, plots and total recovery summary tables were generated with the use of Frapplot package (Guanqiao Ding (2019). frapplot: Automatic Data Processing and Visualization for FRAP. R package version 0.1.3. URL: https://CRAN.R-project.org/package=frapplot).

### CRISPR/Cas9 Knockout of HRI

Oligonucleotides encoding gRNAs targeting the first exon of HRI were designed using CRISPR Design software from the Zhang lab (crispr.mit.edu). Oligonucleotide were annealed and cloned into pCas-Guide (Origene) according to manufacturer’s protocol. gRNA targeting the first exon of HRI contained the following sequence: GCCCTCGGCGGGAAAGTCGA. pCas-guide plasmids were co-transfected with pDonor-D09 (GeneCopoeia), which carries a Puromycin resistance cassette, using Lipofectamine 2000 (Invitrogen). The following day, cells were selected with 1.5 μg/ml of puromycin. Selection was allowed to continue for 24 hrs to lessen the likelihood of genomic incorporation of pDonor-D09. Cells were screened based on their failure to form stress granules or phosphorylate eIF2α after exposure to NaAsO_2_. Cells were cloned by limiting dilution. To confirm genomic ablation of HRI genomic DNA was purified as previously described(Kedersha et al. 2016). Cells were resuspended at 10_8_ cells/ mL in digestion buffer (100 mM NaCl, 10 mM Tris [pH 8.0], 25 mM EDTA [pH 8.0], 0.5% SDS, 0.1 mg/ml proteinase K) and incubated overnight at 55°C. DNA was extracted with phenol/chloroform and precipitated with 2.5 M ammonium acetate and 2 volumes of 100% ethanol, washed with 70% ethanol and air dried. DNA pellet was resuspended in TE containing 0.1% SDS and RNase A (1 μg/ml) and incubated at 37°C for 1 hour. DNA was extracted with phenol/chloroform and precipitated as previously described. Resulting pellet was resuspended at a concentration of 100 ng/μl. For genotyping, the first exon of HRI was amplified by PCR using primers located within the promoter and first intron (Forward: CTAGCTGCAGCATCGGAGT, Reverse: GAGGCAGACGTTCTTTTCAA) using AccuPrime G-C rich polymerase (Invitrogen). Amplicons were cloned into pGEM-T Easy vector (Promega) and sequenced.

## Data Availability

The authors declare that there are no primary datasets and computer codes associated with this study.

## Acknowledgments

We would like to thank Drs. Barbara Sollner-Webb and Denis Lafontaine for helpful comments before submission of this manuscript. This work was funded by grants from the United States NIH (GM124458 to SML, GM126150 to PI, GM126901 to PA) and National Science Centre in Poland (UMO-2015/17/B/NZ7/03043 to WS).

## Author Contributions

S.M.L. conceived, designed and preformed the analysis, interpreted results and took the lead in writing the paper. W.S., M.S., M.L., S.O., A.A., preformed analysis, interpreted results and designed experiments. D.D. and S.M. preformed analysis and aided in sample preparation. P.A. and P.I. gave critical feedback and interpreted results.

---

**Supplemental Figure 1 (A)** Response of U2OS cells to indicated treatments. NaAsO_2_, Lomustine and H_2_O_2_ activate the ISR as indicated by phosphorylation of eIF2α, while H_2_O_2_ also deactivated mTOR as indicated by dephosphorylation of 4EBP1. CHX and puromycin affect neither pathway. (**B**) p-53 null HeLa cells were treated with indicated stressors and rRNA was analyzed by northern blotting. rRNA processing was altered in a similar manner as in p53-positive U2OS cells. (**C**) Untransformed RPE-1 cells were treated with indicated stressors and rRNA was analyzed by northern blotting. rRNA processing was altered in a similar manner as to cancerous U2OS and HeLa cells. **(D)** Mouse NIH3T3 cells were treated with indicated stressors and rRNA was analyzed by northern blotting. rRNA processing was altered in a similar manner as to human U2OS, HeLa and RPE-1 cells. **(E)** Generation of 34S fragment and depletion of downstream precursors occurs in a time- and dose-dependent manner in U2OS cells.

**Supplemental Figure 2** (**A – H**) U2OS cells were treated with NaAsO_2_ and nucleolar markers were used to monitor nucleolar integrity. PHF6 (**A**), RPA194 (**B**), BMS1 (**C**), eIF6, TIF1A (**D**), FBL, NOP16 (**E**), DKC1 (**F**), CIRH1A (**G**), and Ki67 (**H**) were analyzed. This suite of proteins contains representatives of each 3 nucleolar subcompartments. **(I)** Nucleolar integrity is maintained in p53-deficient HeLa cells as monitored by NPM staining.

**Supplemental Figure 3 (A)** Generation of stable cell lines expressing mCherry-NPM, mCherry-Nol9 or EGFP-RPL7A. U2OS cells were transfected with plasmids encoding tagged versions of indicated proteins, drug selected and single cell clones were obtained by limiting dilution. Clones expressing near endogenous levels were selected for future use. Tagged versions are revealed by slower mobility **(B – C)** FRAP of mCherry-NPM reveals decreased mobility of this protein in response to stress in line with inhibition of rRNA processing.

**Supplemental Figure 4 (A)** Genotyping of HRI locus reveals a 40-nucleotide deletion within the 1_st_ exon of HRI. This results in a premature termination codon in the 2_nd_ exon leaving 13 exon junctions between the PTC and the natural termination codon. Red lettering indicates location of gRNA. **(B - C)** Comparison of HRI protein sequence from WT U2OS and hypothetical translation product from ΔHRI cells. Green lettering indicates differences downstream of frame shifting mutation. **(D)** U2OSΔHRI fail to form stress granules in response to NaAsO_2_, but still form stress granules in response to thapsigargin, which activates PERK. Cells were treated with 100 μM NaAsO_2_ or 2 mM thapsigargin for 1 hour and then prepared for immunofluorescence. Cells were stained with DAPI (blue), eIF4G (Green) and eIF3B (Red). **(E)** ΔHRI cells do not inhibit translation in response to NaAsO_2_, but still respond to thapsigargin. U2OS (WT) cells cease translation upon treatment with NaAsO_2_ or thapsigargin. However, U2OSΔHRI cells are refractory to NaAsO_2_-mediated translation inhibition, but maintain their response to thapsigargin. (**F**) Growth curves of U2OS cells and U2OSΔHRI cells demonstrate no statistically significant difference in growth rates between the two cell lines.

